# Identifying and quantifying the disease responses of different pennycress accessions to *Alternaria japonica* and *Sclerotinia sclerotiorum*

**DOI:** 10.1101/2025.05.15.654111

**Authors:** Alice Kujur, Jennette M. Codjoe, Ratan Chopra, Dilip M Shah

## Abstract

Field Pennycress (*Thlaspi arvense*) is gaining attention in the US Midwest as a potential oilseed cover crop for their corn-soybean systems. Field research with breeding trials have shown that pennycress is susceptible to major fungal diseases affecting Brassica crops. In this study, we identified two pathogens: *Alternaria japonica* and *Sclerotinia sclerotiorum* which cause Alternaria black spot and Sclerotinia white mold diseases, respectively. Both fungi infect pennycress leaves and pods, with *S. sclerotiorum* also able to infect stems of the two pennycress accessions tested. We found accession 2032 to be more susceptible than MN106. Traditional visual methods to estimate disease severity could not capture the differential progression of *Alternaria* black spot in the two pennycress accessions. To address this, we developed a cost-effective DNA-based qPCR-based assay that can detect differential growth of both *A. japonica* and *S. sclerotiorum* on leaves and pods of two pennycress accessions. This method also provided more precise quantification of *A. japonica* and *S. sclerotiorum* at early infection stages than was possible visually. This assay could be helpful in evaluating various pennycress cultivars and other crops affected by these pathogens.

## Introduction

*Thlaspi arvense* L., popularly known as stinkweed, fanweed, field pennycress, or simply pennycress, is an annual winter weed with roots tracing back to Eurasia. It was introduced in the U.S. around the 1700s. It has since adapted impressively to diverse environments, thriving in disturbed non-agricultural areas and agricultural lands like pastures and croplands across various soil types. This remarkable plant belongs to the mustard family (Brassicaceae). It is closely related to valuable oilseed crops, such as rapeseed (*Brassica rapa* and *Brassica napus L.*) and camelina (*Camelina sativa L.*), alongside the model plant *Arabidopsis thaliana*. The potential of pennycress to enhance ecosystems and boost agricultural productivity while minimizing land use is revolutionary (Keadle et al., 2022; Basnet and Ellison, 2023). Its seeds not only yield oil suitable for low-carbon biofuels—such as renewable biodiesel and sustainable aviation fuel—but also serve as a high-protein animal feed that is crucial in today’s food systems.

Ecologically, pennycress plays a vital role in sequestering carbon and preventing organic soil carbon loss through effective cover cropping. It contributes to greenhouse gas reduction, improves soil health (including pest control for threats like the Fall armyworm), mitigates soil erosion, and reduces nutrient runoff— making it a powerful ally in sustainable farming practices. Current initiatives aim to domesticate, breed, and commercialize pennycress as a high-value oilseed rotational cash cover crop, particularly in the Midwestern United States, where it can flourish on fallow lands currently used for corn-soybean rotations (Sedbrook et al., 2014). However, pennycress can face challenges from various pathogenic fungi that commonly affect other Brassicaceae crops (Keadle et al., 2022; Basnet and Ellison, 2023). Alternaria leaf spot (ALS) and Sclerotinia white mold (SWM) stand out as the two most damaging fungal diseases, presenting a significant challenge to the future of pennycress cultivation.

ALS encompasses a spectrum of fungal diseases that can target a wide variety of plant hosts beyond Brassica species. Known under various aliases—such as Alternaria black spot (ABS), leaf blight, sooty spot, and pod spot—these diseases are caused by diverse *Alternaria* species, which are often dark-pigmented and can infect over 4,000 plant species (Meena et al., 2017; Wodenberg et al., 2015). With more than 20% of agricultural yield losses attributed annually to *Alternaria* species, it is critical to recognize potential threats posed by *A. brassicae*, *A. brassicicola*, and *A. raphani*, which can reduce seed yields exceeding 80%. These necrotrophic ascomycete pathogens are particularly severe during flowering and ripening stages, making timely intervention essential. Additionally, SWM represents another formidable adversary to crop health, caused by three *Sclerotinia* species *S. sclerotiorum*, *S. minor*, and *S. trifolium*. These fungi can wreak havoc on over 370 plant species across 64 families, leading to yield losses of up to 80% (Li et al., 2006). The disease is marked by necrotic brown lesions that transform into a distinctive fluffy white mycelial growth—an alarming sign that sclerotia, the compact resting bodies, are forming. Sclerotia can persist in the soil for years, making the disease difficult to eradicate.

Given that pennycress can be susceptible to these pathogens, the risk of increased disease outbreaks looms ever larger as pennycress cultivation intensifies. If not properly managed, the impact on yield, seed quality, and oil content could be devastating. The varying responses of different fungal species to fungicides underscore the need for proper taxonomic identification and characterization of these pathogens. Therefore, proactive isolation along with morphological and molecular identification of these disease-causing fungi is vital.

In this study, we describe techniques to identify *Alternaria* and *Sclerotinia* species infecting pennycress and developed disease assay protocols to screen for resistance in controlled conditions. All pathogenicity assays were developed using two natural accessions of pennycress, MN106 and 2032, with different degrees of biotic stress tolerance. MN106 served as a line with wild-type levels of disease resistance. In contrast, 2032 plants often succumbed to unknown diseases in CoverCress Inc. field and greenhouse experiments and served as a susceptible line in our experiments.

## Materials and Methods

### Fungal isolates collection, culturing, and preservation

Field-grown pennycress pods exhibiting symptoms of ABS disease were collected in 2021 from four different locations in the United States: Manito, IL; Sigel, IL; Venedy, IL; and Mt. Pulaski, IL. These symptomatic pods were used as primary inoculum to culture *Alternaria* species on Petri plates containing Potato Dextrose Agar (PDA; Fisher Scientific, Pittsburgh, PA) supplemented with antibiotics (0.02 mg/ml each of chloramphenicol and streptomycin sulfate). The plates were incubated at 25°C in the dark for 5 days. After this incubation, the primary cultures were sub-cultured onto fresh PDA plates with the same media and growth conditions.

After 8 days, about 4-5 mycelial discs (8 mm in diameter) from the subcultured PDA plates were stored in 30% glycerol solution or in Potato Dextrose Broth (PDB) medium with 0.02 mg/ml of chloramphenicol and streptomycin sulfate, and 30% glycerol at -80°C for long-term storage. The 8-day-old subculture plates were also utilized to initiate subsequent cultures. These cultures were maintained through subculturing by transferring small mycelial discs from the growing edges of 14-day-old cultures (using a sterile stainless-steel blade) to the centers of fresh PDA plates containing antibiotics. A new culture was established every 6 weeks from the mycelial discs stored in the 30% glycerol solution at -80°C.

For culturing *Sclerotinia* species in the laboratory, sclerotia were collected from pennycress plants exhibiting white mold disease in a field in Sigel, IL, USA, in 2021. A single sclerotiorum was sterilized using 50% bleach, then cut open and incubated on a water agar plate. The mycelium that grew from the sclerotiorum was transferred to a PDA plate and incubated at 25°C in the dark. This plate served as the basis for subsequent cultures. Subculturing of *Sclerotinia* was done every three days by taking a mycelial punch (using the back of a sterile P200 pipette tip) from one PDA plate to transfer to a fresh PDA plate. A new culture was also initiated from a sclerotiorum from the master plate every 6 weeks.

### Fungal genomic DNA extraction, PCR amplification and sequencing

We isolated genomic DNA (gDNA) from the *Sclerotinia* field isolate from 3 ml of liquid culture grown in PDB (protocol described below) for 2 days. For *Alternaria* field isolates, we extracted gDNA from about 100 mg of 30-day-old mycelia scraped from the flat edges of PDA plates with 0.02 mg/ml of antibiotics. We followed the E.Z.N.A.® Fungal DNA Mini Kit (Omega, Bio-Tek, USA) instructions for the gDNA extraction. To amplify the Internal Transcribed Spacer (ITS) region of the 5.8S rRNA gene from both *Alternaria* and *Sclerotinia* isolates, we used universal forward, and reverse primer sets from earlier studies (ITS1 and ITS4 from Meena et al., 2017 for *Alternaria*, and ITS4 and ITS5 from Baturo-Ciesniewska et al., 2017 for *Sclerotinia*).

For PCR amplification, we used about 60 ng of gDNA from the *Alternaria* isolates and 250 ng from the *Sclerotinia* isolate. A reaction mix was prepared containing Q5® High-Fidelity DNA Polymerase (New England Biolabs, Inc., USA), 5X Q5 Reaction Buffer, 10 mM dNTPs, and 10 µM of each primer, totaling to 50 µl per sample. The cycling conditions were performed using a BioRad thermal cycler by setting the initial 30 seconds of denaturation at 98°C, followed by 35 cycles of 10 seconds of denaturation at 98°C, 15 seconds of annealing at 63°C, and 30 seconds of elongation at 72°C. Finally, the process was completed with a 2-minute elongation at 72°C. The sizes of the amplicons were compared on a 1% agarose gel in 1X TAE buffer. The amplicons from the *Alternaria* and *Sclerotinia* isolates were purified using the QIAquick PCR Purification Kit (QIAGEN, Germany) and the Monarch DNA Gel Extraction Kit (NEB), respectively, and then sequenced using the universal ITS primers mentioned above. Finally, the nucleotide sequences were aligned in Benchling to get consensus sequences for each of the *Alternaria* and *Sclerotinia* isolates. A BLASTN query was performed against similar sequences in the NCBI GenBank database.

### Plant materials and growth conditions

In this study, two natural pennycress accessions from North America, namely MN106 and 2032, were used for all pathogenicity tests. The seeds were obtained from CoverCress™ Inc. in St. Louis, MO, USA. Seeds were soaked in a solution of 3.3 µg/ml gibberellic acid 4+7 overnight before sowing them in Pro-Mix FPX with a fungicide mix. After two weeks, seedlings were transplanted into pots with a Berger BM7 35% bark mix. We grew MN106 and 2032 plants in a growth chamber with 14 hours of light at 25°C with a light intensity of 300 µmol followed by 10 hours of darkness at 23°C and 50% relative humidity. For pod and stem assays, 2-week-old seedlings were vernalized for 3 weeks in a 4°C cold room then returned to growth chambers.

### *A. japonica* pathogenicity assay

A preliminary pathogenicity assay was conducted to evaluate field isolates of *A. japonica* from Manito, IL, and Sigel, IL, using leaves from non-vernalized, 4.5 to 5-week-old MN106 and 2032 plants at the vegetative stage. Ten healthy plants of each accession and a two-week-old *A. japonica* subculture were used for the assay. Six mature rosette leaves (the 7^th^ to 12^th^ leaves in ascending order) from each plant were detached and placed on soaked paper towels with 3 ml of sterile water in Petri plates. Mycelial agar plugs, measuring 2 mm in diameter, were taken from the growing edge of *A. japonica* on PDA and placed on each leaf from five of the plants for inoculation. Each leaf received one mycelial plug with the mycelium side down on the adaxial surface of the wider leaf lamina, while avoiding the veins. Additionally, six leaves from five other plants were mock-inoculated with clean agar plugs to serve as negative controls. The Petri plates containing the mock- and pathogen-inoculated leaves were covered with lids and incubated within airtight containers (plastic storage totes) lined with wet paper towels to maintain high humidity at room temperature in the dark for at least five days.

Each leaf was then examined for the infected lesion area. The assay with the *A. japonica* field isolates from Sigel, IL was repeated at least for four times.

Furthermore, a pathogenicity assay of *A. japonica* on pods was conducted using the field isolate from Sigel, IL, on both green mature and yellow dried pods collected from vernalized, healthy plants of each accession. Six green mature pods and six yellow dried pods were randomly detached from the main stem of eight plants of each accession in ascending order. The green mature pods were taken from approximately 12-week-old plants, while the yellow dried pods were taken from plants older than 12.5 weeks. Subsequently, as in the previous assay, pods from four plants were inoculated with *A. japonica* mycelial agar plugs measuring 2 mm in diameter, while pods from the other four plants were mock-inoculated with clean agar plugs to serve as negative controls. Inoculated pods were subjected to the same conditions as the inoculated leaves and observed for five days. This assay was also conducted at least three times.

### *S. sclerotiorum* pathogenicity assay

A pre-screening infection assay for *S. sclerotiorum* was conducted on the leaves of non-vernalized plants aged 4.5 to 6 weeks, specifically MN106 and 2032, at the vegetative stage, following the methodology established for the pathogenicity assays for *A. japonica*. Nine mature rosette leaves, specifically the 5th to 13th leaves, were harvested from five plants per accession and subsequently placed on Petri dishes lined with paper towels that had been saturated with 2 ml of sterile water. The adaxial surface of the leaves, avoiding the veins, were inoculated with 2 mm diameter agar plugs containing *S. sclerotiorum* mycelium (3 days after transfer to fresh PDA plate) by positioning the mycelium-side down. These inoculated leaves were maintained under conditions identical to those applied to the inoculated leaves and pods for the *A. japonica* pathogenicity assays.

Subsequent visual pathogenicity assays were conducted on the 7^th^, 8^th^, and 9^th^ rosette leaves. At one-, two-, and three-days post-inoculation (dpi), the necrotic lesion area was quantified using FIJI software based on images captured with a camera or via CropReporter (PhenoVation, Wageningen, Netherlands).

To test pathogenicity on pods, a suspension comprising fragmented mycelium from liquid cultures of *S. sclerotiorum* was employed as the inoculum, following the protocol adapted from Djami-Tchatchou et al. (2023). Mycelium was scraped from an area of approximately 2 cm² on 4 to 5-day-old PDA subculture plates, homogenized in 1 ml of PDB using a plastic pestle within a microcentrifuge tube. A volume of 0.3 ml of this fragmented mycelium was inoculated into 3 ml of PDB and incubated at 28°C with shaking at 220 rpm for 2 days. The mycelium obtained from the 3 ml of PDB culture was filtered using a 70 µm cell strainer and transferred to a microcentrifuge tube containing 1 ml of ½X PDB for homogenization using a plastic pestle. The concentration of the fragmented mycelium was subsequently adjusted to an optical density at 600 nm of 0.5 with ½X PDB, and the suspension was applied to detached pods using a refillable 3 ml perfume atomizer. The inoculated pods were incubated in a sealed plastic tote at high relative humidity, at room temperature, and in the dark for a duration of 4 to 5 days until symptoms manifested. Twelve pods, including those of varying ages, were harvested from the main stem to facilitate this assay.

Furthermore, pennycress stems were also subjected to inoculation with *S. sclerotiorum*. Inoculation of the stems were performed by excising the primary inflorescences just below the main meristem and employing mycelial agar plugs from the leading edge of a 3-day-old *S. sclerotiorum* PDA subculture, following the methodology outlined in Koga et al. (2014). The inoculated plants were maintained in a growth chamber at 95% humidity for 9 days, during which lesion lengths were recorded every other day.

### DNA-based real-time quantitative PCR primers designing

We designed specific forward and reverse primer pairs derived from the DNA sequences of the internal transcribed spacer (ITS) region of the 5.8S ribosomal RNA (rRNA) genes of *A. japonica* and *S. sclerotiorum*. Additionally, we too designed primer pairs targeting an essential housekeeping gene, *TaActin* (Thalr.0014s0459), found in the pennycress (*T. arvense*) genome. The design process was executed using BatchPrimer3 v2.0 software, which is tailored for creating primers with high specificity. While all other parameters were maintained at their default settings, we carefully adjusted the product size range (100-200 bp) to meet our experimental requirements.

### Sample collection, genomic DNA isolation, and DNA-based real-time quantitative PCR assay

The pathogenicity assays for *A. japonica* and *S. sclerotiorum* described above were repeated on detached 8^th^ rosette leaves and pods from the 2032 and MN106 lines. Leaf discs, each 22 mm in diameter, were collected by removing plugs from mock and pathogen-inoculated leaves at various time points. These included samples taken at 3-6 dpi for the *A. japonica*-based assays and at 1-3 dpi for the *S. sclerotiorum*-based experiments. The discs collected from pathogen-inoculated leaves encompassed visible lesions. Three leaf discs, one from each leaf, were combined to create a single biological replicate for each treatment at each time point. A total of three biological replicates per treatment at different time points for each accession were collected, flash-frozen in liquid nitrogen, and stored at - 80°C before processing.

Similarly, pods from randomly selected plants of the 2032 and MN106 lines were mock- and pathogen-inoculated. Three pods, each from a different plant of each accession, were pooled to form one biological replicate for each treatment at each time point. The pod samples were collected after removing the plug at 2-5 dpi for the *A. japonica*-based assays. Whereas, for the *S. sclerotiorum*-based experiments, inoculum sprayed pods at 1-4 dpi and uninoculated pods at 0 dpi were collected. As with the leaves, three biological replicates of pod samples were flash-frozen in liquid nitrogen and stored at -80°C before processing.

gDNA was isolated from every biological replicate sample of leaves and pods using the Quick-DNA Plant/Seed Miniprep Kit (Zymo Research, USA), following the manufacturer’s instructions. At least two technical replicates per biological replicate were included in the gDNA isolation. Samples were homogenized in safe-lock microcentrifuge tubes using a TissueLyser (QIAGEN, Hilden, Germany) for two sets of 5 minutes at a frequency of 25 oscillations per second. For DNA-based qPCR assays, gDNA was diluted to 25 ng/µl to ensure that the final qPCR mixture contained 5 ng of gDNA as a template. A total volume of 10 µl of the qPCR reaction mix was prepared, consisting of 5 µM each of forward and reverse gene-specific primers (ITS primers for the pathogen and *TaActin* primers for pennycress) and 5 µl of SYBR Green qPCR 2X Master Mix (Intact Genomics, USA). The qPCR was performed in triplicate for each biological replicate using the Biorad CFX Real-Time System. The cycling conditions included an initial step at 95°C for 15 minutes, followed by 40 cycles of denaturation at 95°C for 5 seconds, annealing at 60°C for 30 seconds, final elongation at 65°C for 5 seconds, and melting curve analysis at 95°C for 5 seconds. A negative control and no-template control were included for each primer pair. The results were analyzed using the CFX Real-Time System software and Microsoft Office Excel based on the obtained cycle quantification (Cq) values. The relative abundance of the ITS-5.8S rRNA gene to *TaActin* was calculated as 2^-ΔΔCq^. A representative result was presented as the mean 2^-ΔΔCq^ value ± SEM from sets of results derived from three biological replicates (each consisting of technical triplicates) for each treatment at different time points for each accession.

### Trypan blue staining

The process of Trypan blue staining was conducted on both mock-inoculated and *A. japonica*-inoculated leaves (specifically the 9th rosette leaves) as well as pods from the 2032 and MN106 plants. Following the established methodology outlined by McDowell et al. (2011), the staining was performed at several key time points. For the leaves, samples were collected and stained from 3 to 6 dpi, while for the pods, the staining was carried out from 2 to 5 dpi. Stained lesions were quantified from camera images using FIJI. Trypan blue staining was performed more than three times.

### Data analysis

The qPCR assay results were analyzed and visualized using Microsoft Office Excel. For statistical analysis, we relied on the Pandas, a Python library used for data manipulation and analysis. To determine significant differences between the sample groups, we applied Welch’s t-test, which is particularly suitable for comparing means when the variances of the two groups are unequal.

## Results

### Fungal disease symptoms and pathogen isolation from field-grown pennycress

Severe fungal disease symptoms were observed on field-grown pennycress at several locations in the U.S., including Manito, Sigel, Venedy, and Mt. Pulaski, IL, during research trials by CoverCress™ Inc. in 2021. These diseases affected the leaves, stems, roots, flowers, pods, and seeds of pennycress. Seeds from these plants were analyzed by the Seed Testing Laboratory at Iowa State University, revealing multiple fungal pathogens, especially those causing ALS, pod black spot, and SWM. To culture these pathogens for further characterization, we used pennycress pods with *Alternaria* black spot symptoms and sclerotia from plants with SWM as inoculum for isolation. Fungal isolates, particularly *Alternaria* spp., were initially characterized by their macroscopic features. The *Alternaria* isolates from Manito and Sigel, IL, showed a cottony olive-green color on PDA plates after 8 days of incubation at room temperature (RT) in the dark. The *Sclerotinia* isolate appeared white on PDA plates after 3 days at the same condition.

Efforts to induce sporulation using various methods were unsuccessful. As a result, we purified the isolates by subculturing the growing edges of the mycelia onto new PDA plates.

### Molecular identification of. *A. japonica* and *S. sclerotiorum* in pennycress

The ITS regions of 560 and 599 base pairs from 5.8S rRNA genes were successfully amplified and sequenced from *Sclerotinia* and *Alternaria* field isolates using fungal-specific universal ITS primer pairs, respectively. The sequences for the forward and reverse ITS primer pairs used in this study are provided in Table 1. A BLASTN analysis of the sequenced ITS amplicon from the 5.8S rRNA gene of the two *Alternaria* field isolates, collected from Manito and Sigel, IL, revealed similarities of 97.78% and 96.57% to the sequences of *A. japonica* with accession numbers MK940392.1 and KY788039.1, respectively, in the NCBI GenBank database. The ITS-5.8S rRNA amplicon sequence from the *Sclerotinia* field isolate showed 100% identity with numerous ITS sequences of *S. sclerotiorum* strains, including accession numbers MN216247.1 and MH318571.1. The top 10 BLAST hits for *A. japonica* and S. *sclerotiorum* can be found in **Supplemental Fig 1**. The ITS-5.8rRNA sequences isolated from *A. japonica* strain Sigel, *A. japonica* strain Manito, and the *S. sclerotiorum* strain Sigel have been submitted to NCBI-GenBank database with accession numbers PV626842-PV626844.

**Table 1:**
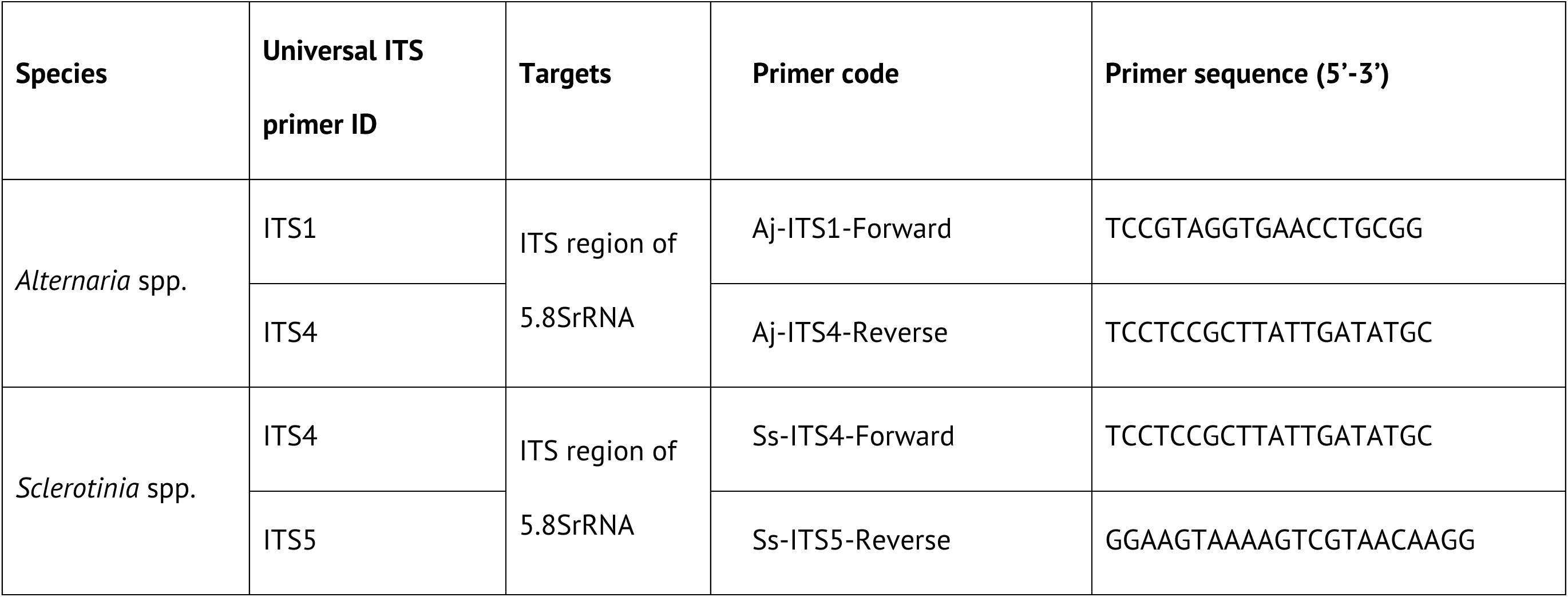
Details of primers used in this study for sequencing the isolates of *A. japonica* and *S. sclerotiorum* from pennycress field.

### Pathogenicity of *A. japonica* on leaves and pods

A preliminary infection assay with detached leaves allowed for quick screening of virulent *A. japonica* field isolates for pathogenicity in pennycress. Leaves from MN106 and 2032 plants were inoculated with *A. japonica* mycelial agar plugs or mock-inoculated with clean PDA plugs and examined for disease symptoms up to 5 dpi. Both field isolates from Manito, IL, and Sigel, IL, showed disease symptoms in pennycress at 3 dpi. However, the subsequent assays focused on the isolate from Sigel, IL, due to discoloration observed in the Manito isolate during sub-culturing. By 4 dpi, all *A. japonica*-inoculated leaves showed distinct concentric sunken lesions of fungal biomass, while mock-inoculated leaves remained symptom-free (**Fig 1a and 4**). There was no noticeable difference in lesion size between the two leaf types or among different leaf ages. A prominent yellow halo distinguishable from age-related leaf senescence was however seen around lesions on MN106 leaves. We suspected higher fungal biomass in 2032 leaves although this could not be quantified from images. Trypan blue staining of infected leaves showed larger lesions on 2032 leaves compared to MN106 leaves (**Fig 1b**), which were further measured and found to be significantly larger on 2032 leaves (**Fig 1c)**. The Trypan blue-stained lesions likely consist of both fungal hypha and dead plant cells.

**Fig 1:**
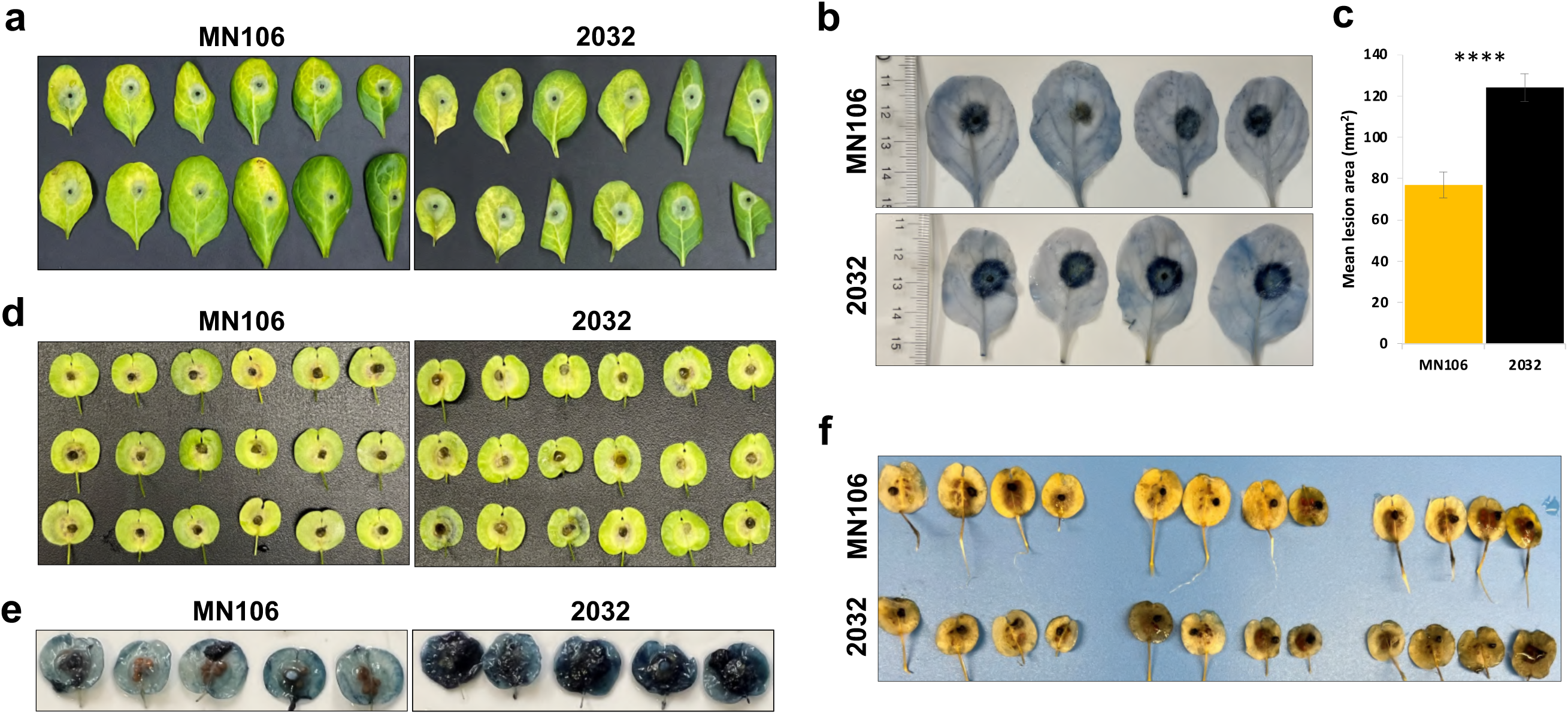
The pathogenicity assay of *A. japonica* demonstrated its ability to infect pennycress leaves and pods of all ages. (a) Representative images show *A. japonica*-infected leaves from 7th to 12th positions (left to right) of two pennycress accessions, MN106 (left) and 2032 (right), at 4 days post-inoculation (dpi). The leaves were detached from non-vernalized, 4.5-week-old plants and inoculated with 2 mm *A. japonica* mycelial plugs. (b) This panel displays hyphal growth and cell death in the *A. japonica*-infected 8th to 11th leaves of MN106 (top row) and 2032 (bottom row), stained with Trypan blue at 4 dpi. (c) This panel shows the measurement of the relative mean lesion area (mm²) in the leaves of MN106 and 2032 after Trypan blue staining at 4 dpi. Each data point represents the mean lesion area from four 9th leaves per plant. The data presented are pooled results from two separate experiments. An indication of **** represents a p-value < 0.0005 based on the two-tailed t-test assuming unequal variances. (d-e) These representative pictures illustrate *A. japonica*-infected mature green pods of the two pennycress accessions, MN106 (left) and 2032 (right) at 4 dpi. Panel (d) shows the pods without Trypan blue staining, while panel (e) shows them with the staining. The pods were detached from vernalized 12-week-old plants and inoculated with 2 mm *A. japonica* mycelial plugs. (f) This panel reveals the dense, darker grey, sooty mycelium accumulated on the dried yellow pods of 2032 (bottom row) compared to those of MN106 (top row).

In assays on detached green mature pods, 2032 pods appeared to accumulate more fungal biomass than MN106 pods at 3 dpi, although that is not evident from images (**Fig 1d)**. Trypan blue staining confirmed more abundant hyphal growth and cell death in 2032 pods than in MN106 pods at 4 dpi, but not in a way that was easily quantifiable (**Fig 1e**). When dried yellow pods were inoculated with *A. japonica,* 2032 pods had a dense coating of darker grey mycelium compared to MN106 pods (**Fig 1f**).

Additionally, we monitored the lesion development through Trypan blue staining over time (3-6 dpi) on *A. japonica-*inoculated leaves of both lines to check if symptoms varied across different stages of plant-fungus interaction. The lesion diameter increased over time in both genotypes. 2032 leaves had larger lesions than MN106 at all time points, although significantly different only for days 3-5dpi (**Fig 2a-b**).

**Fig 2:**
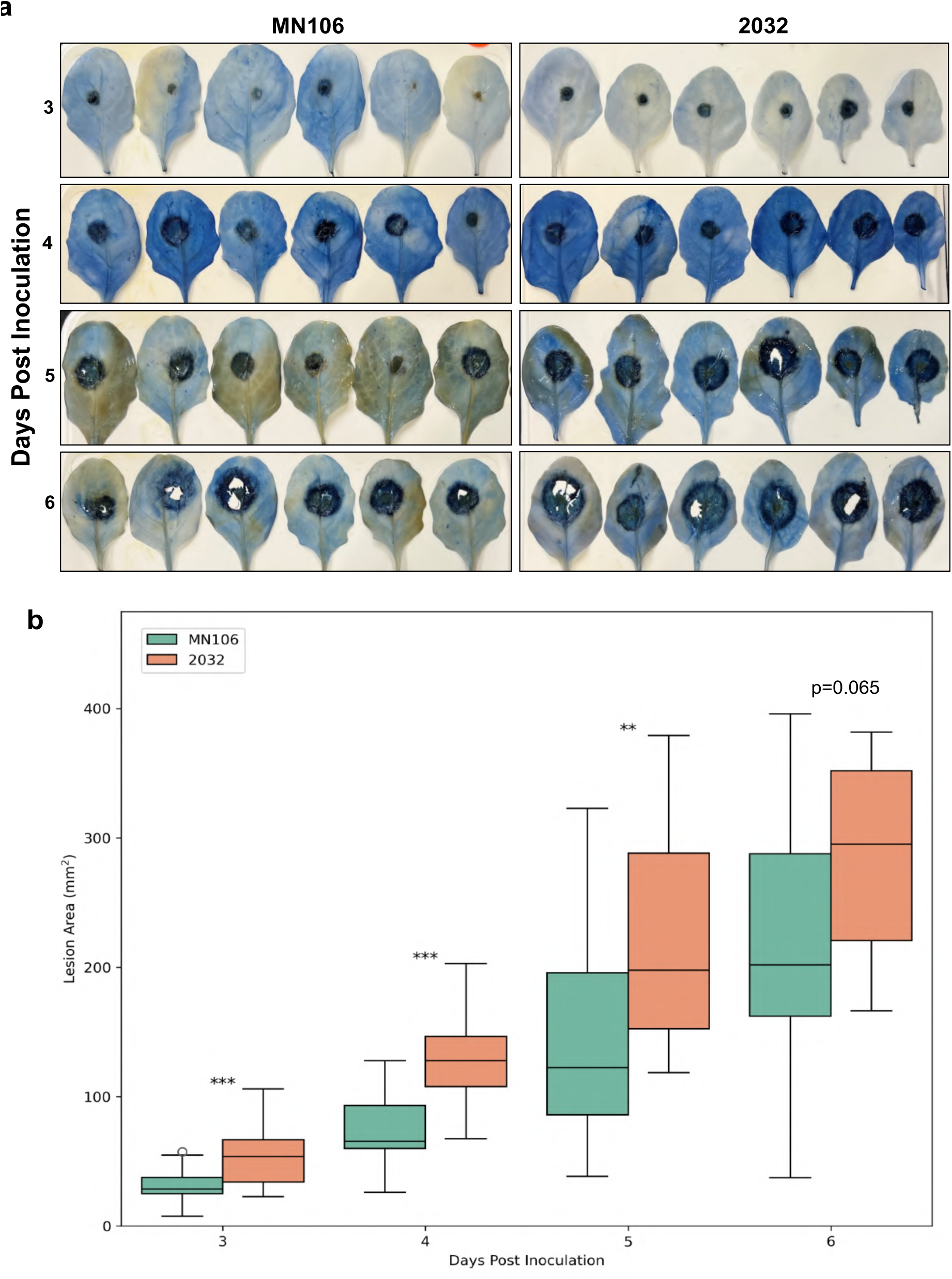
Time-course analysis of *A. japonica* growth on infected leaves of MN106 and 2032 plants. a) The 9th leaves of non-vernalized, 4.5-week-old plants were detached, inoculated with *A. japonica* plugs, incubated for a 3-6 dpi, and stained with Trypan blue. b) The boxplot shows the necrotic lesion area measured from days 3 to 6 dpi in two pennycress accessions. Plotted is the mean necrotic lesion area (mm²) derived from up to six independent experiments. Asterisks indicate statistical significance, with *** representing p-values ≤ 0.0005 and ** indicating p-values ≤ 0.005, as determined by Welch’s t-test.

### Pathogenicity of *S. sclerotiorum* on leaves, pods, and stems

A similar detached leaf pathogenicity assay of *S. sclerotiorum* was undertaken using leaves of different ages. *S. sclerotiorum* inoculation led to the development of brown, necrotic lesions (**Fig 3)**. Depending on the leaf age, these lesions were larger in 2032 compared to MN106 leaves (**Fig 3a**). Interestingly, the 8^th^ rosette leaf showed a clear difference in susceptibility regardless of plant age (**Fig 3a**).

**Fig 3.**
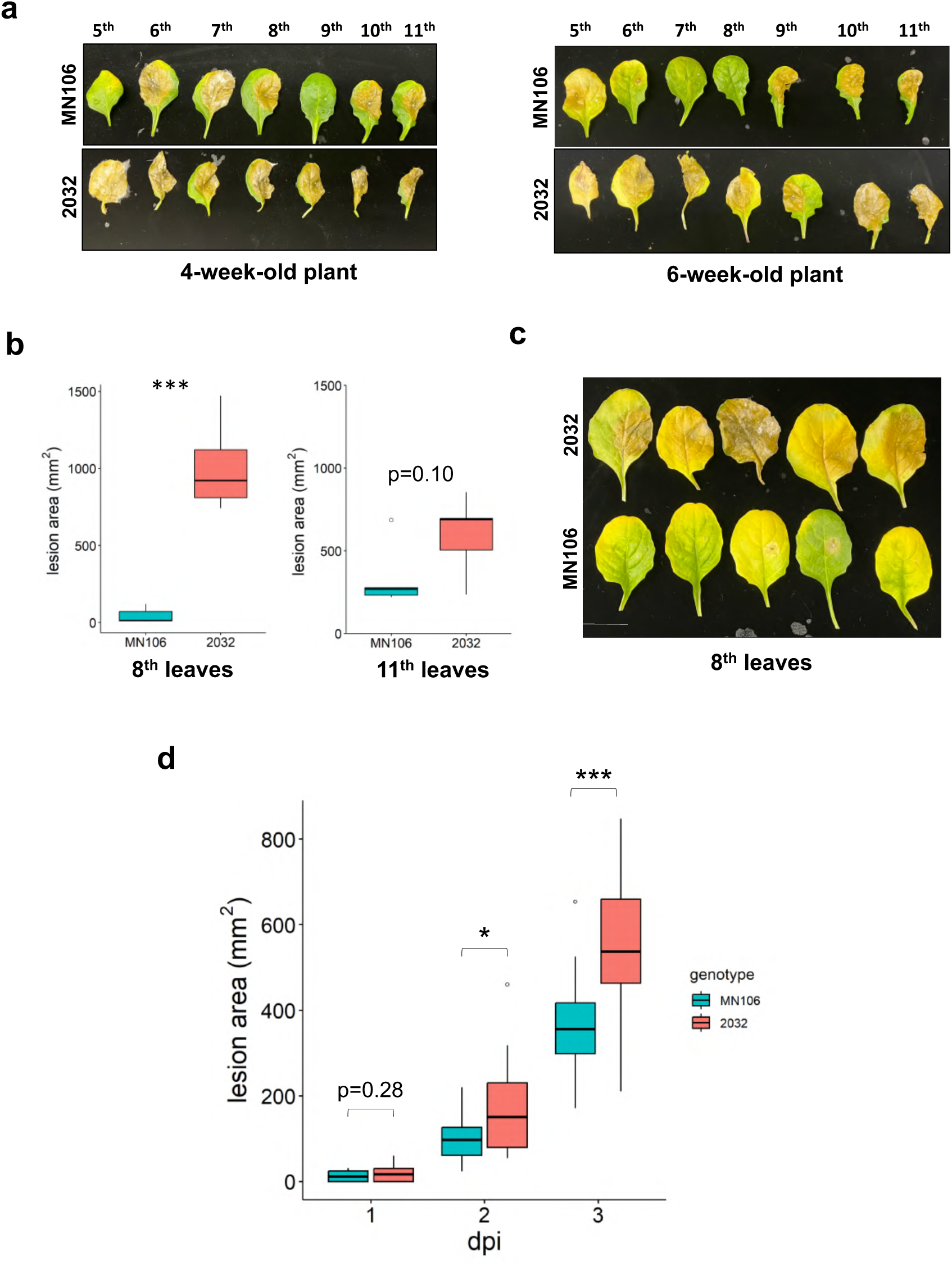
Demonstration of pathogenicity of *S. sclerotiorum* on the leaves of twopennycress accessions, 2032 and MN106 with different susceptibility. Panel (a) shows photographs of the infected leaves (5th to 11th leaves from 4-week-old and 6-week-old plants, respectively) at 3 dpi. Panel (b) depicts necrotic lesion area at 3 dpi of n=5 plants per genotype on the 8th and 11th rosette leaves of 6-week-old plants. (c) Image of the 8th leaves quantified in (b). Scale = 3 cm. d) The 7th, 8th, and 9th leaves of 5-week-old plants were infected with *S. sclerotiorum* agar plugs and the average lesion area on those 3 leaves was calculated for each plant (n=21 plants per genotype across 3 separate experiments) at 1, 2, and 3 dpi. Pooled are the results of 3 separate experiments: (n=21 plants per genotype). (b, d) Statistical significance was determined using Student’s t-test. * indicates p <0.05, *** indicates p <0.001.

**Fig 4.**
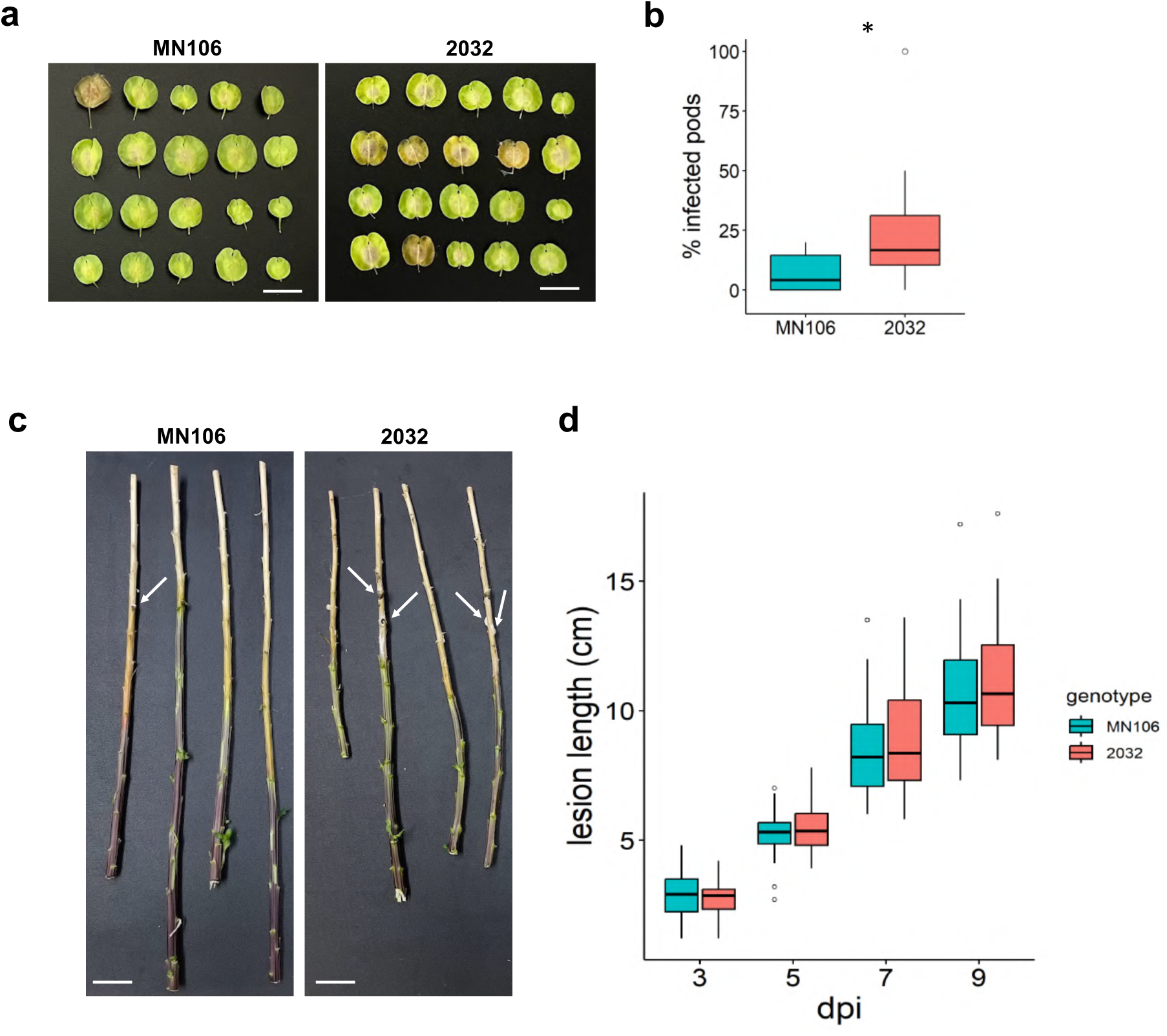
Illustration of higher susceptibility of 2032 pods and stems than those from MN106 plants to *S. sclerotiorum*. Panel (a) shows pods at 5 dpi that were spray-inoculated with *S. sclerotiorum* fragmented mycelia. Each row of pods came from a stems at 9 dpi. Arrows indicate the location of sclerotia. Scale = 2 cm. (d) The length of necrotic lesions that spread down the stem were measured over time from 3 to 9 dpi. Presented is pooled data from 3 separate experiments: n=26 plants per genotype. Pairwise comparisons of the two genotypes at each dpi by Student’s t-tests found no significant differences.

Lesion size quantification on the 8^th^ leaves of five 6-week-old plants per genotype revealed significantly larger lesions on 2032 leaves than MN106 leaves at 3 dpi (**Fig 3b, c**) as compared to the less pronounced difference in susceptibility when using younger 11^th^ leaves (**Fig 3b**). This prompted us to focus on the 8^th^ leaves and those of similar aged leaves for further assays.

The time-course pathogenicity assays of *S. sclerotiorum* utilized the 7th, 8th, and 9th leaves, and an average lesion size from these three leaves per plant was calculated to minimize leaf-to-leaf variations in susceptibility. The results of four individual experiments (using 6-8 plants per genotype each) can be viewed in **Supplemental Fig 2**. The pooled results from three of those experiments are visualized in **Fig 3d**. Looking at the pooled data, at 1 dpi, there was no appreciable difference in lesion area on the two genotypes, but by 2 dpi and especially by 3 dpi, there were significantly larger lesions on 2032 leaves as compared to MN106 leaves (**Fig 3d**).

The same infection assay on pods using mycelial plugs of *S. sclerotiorum* caused such rapid, severe infection that it was impossible to observe the differences in pathogenicity between the two accessions (data not shown). Therefore, detached pods were spray-inoculated with fragmented *S. sclerotiorum* mycelia at an optical density (OD) 600 nm of 0.5. On average, 25% of 2032 pods developed necrosis compared to 7% for MN106 across multiple experiments (**Fig 4a-b**). The spread of necrosis on cut stems was similar for both accessions over time (3-9 dpi) (**Fig 4-d**), although more sclerotia formed on the 2032 stems than on the MN106 stems as observed at 9 dpi (**Fig 4c**).

### Development and validation of qPCR primers for quantification of *A. japonica*, *S. sclerotiorum,* and pennycress DNA

We wondered if qPCR-based assays to detect fungal DNA as a proxy for fungal biomass might be more sensitive at detecting differences in MN106 and 2032 susceptibility to *A. japonica* and *S. sclerotiorum* than the visual assays described above. We designed primers to specifically amplify the ITS region of the 5.8S rRNA genes of either *S. sclerotiorum* or *A. japonica*, as well as a primer pair to amplify a pennycress housekeeping gene, *TaActin* (Thalr.0014s0459) (**Table 2**). We confirmed the specificity of the primers by setting up PCRs for gel-based analysis. gDNA from mock- and pathogen-inoculated leaves and pods at 3 dpi served as templates. Controls were set up with *A. japonica* strain gDNA used as a positive control for Aj-ITS-qPCR primers and a negative control for Ss-ITS-qPCR primers, and vice versa for *S. sclerotiorum* gDNA. The *TaActin* primers amplified a product only from mock- and infected pennycress DNA samples, but not from fungus-only samples or no-template controls (**Fig 5a**). The Aj-ITS-qPCR primers amplified a product from *A. japonica* gDNA and *A. japonica-*inoculated leaf samples, but not from mock-inoculated leaves, *S. sclerotiorum-*inoculated leaves, and *S. sclerotiorum* gDNA (**Fig 5b**). Likewise, the Ss-ITS-qPCR primers amplified *S. sclerotiorum-* containing samples only (**Fig 5c**). Dissociation curve analysis on qPCR amplicons from all tested DNA samples showed that each selected primer pair produced a single peak, re-confirming their specificity (**Fig 5d-e**).

**Fig 5:**
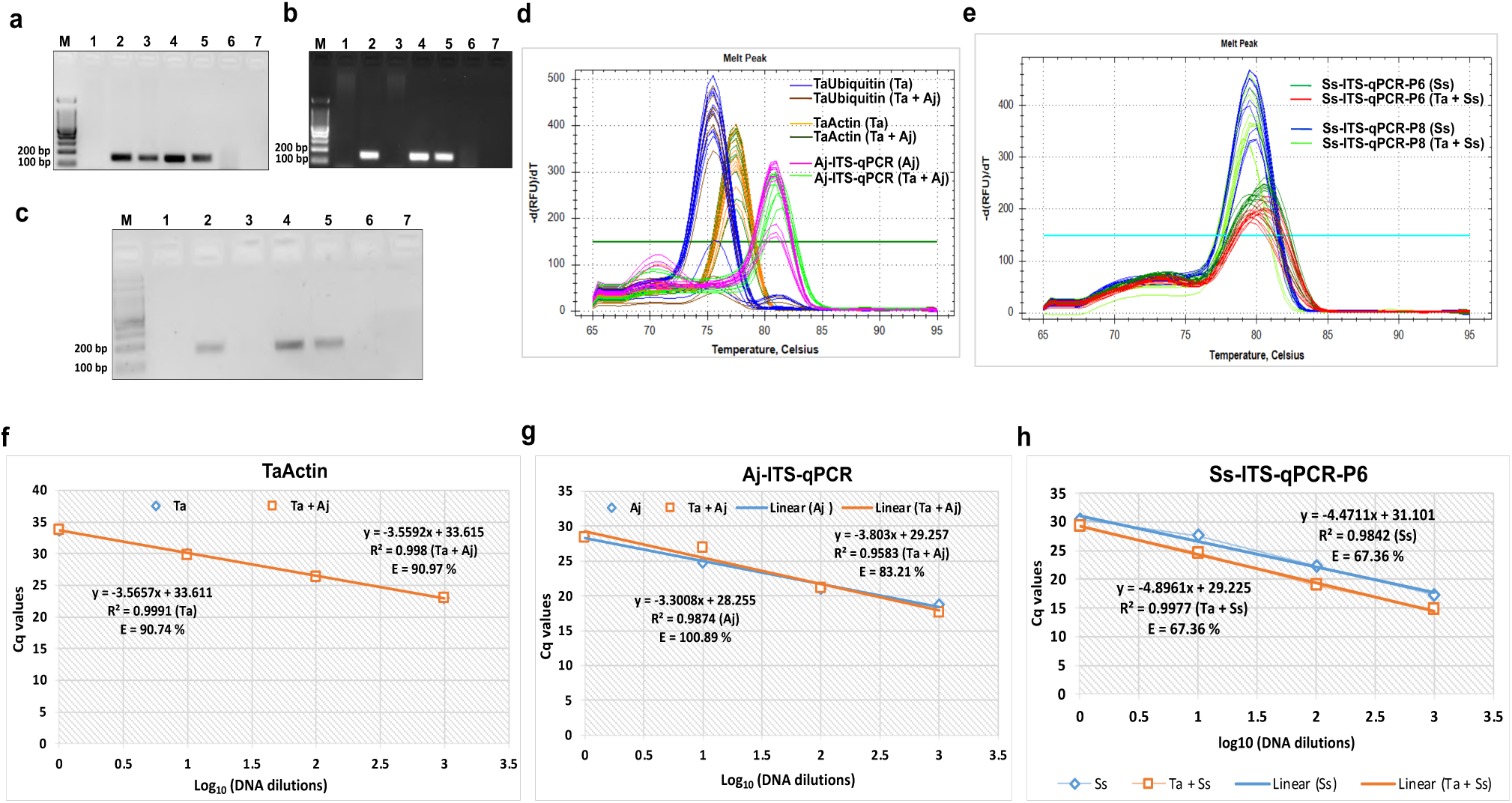
Validation of real-time PCR primers for biomass quantification of *A. japonica*, *S.* two pennycress accessions at 4 dpi. (a) The gel image shows the amplified portion of the *Ta*Actin gene. The marker lane (M) contains a 100 bp Plus DNA Ladder. Lanes 1, 6, and 7 contain gDNA from *A. japonica*, *S. sclerotiorum* and a no-template control, respectively. Lanes 2 and 3 represent mock-inoculated and *A. japonica*-infected samples from MN106; Lanes 4 and 5 shows mock-inoculated and *A. japonica*-infected samples from the 2032 accession. Gel image (b) and (c) display the single amplicons of the Aj-ITS-5.8SrRNA and Ss-ITS-5.8SrRNA genes, respectively. Lanes 1 and 2 include gDNA from mock-inoculated and pathogen-infected samples from MN106, while lanes 3 and 4 represent mock-inoculated and pathogen-infected samples from the 2032 accession in both the images (b) and (c). Lanes 5 and 6 contains gDNA from *A. japonica* and *S. sclerotiorum* in (b) and vice-versa in (c), lane 7 in (b) and (c) is a no-template control. (d-e) The dissociation curve analyses of all target gene amplicons from qPCR using gDNA of uninfected pennycress leaves (Ta) and *A. japonica* (Aj), *S. sclerotiorum* (Ss), and pathogen-infected pennycress plants (Ta + Aj and Ta + Ss) were performed. The qPCR products from each validated primer pair exhibited a single melt curve above the threshold value, indicating the denaturation of only one PCR product for each primer pair. (f-h) The percentage amplification efficiency (E) of the validated primers for gene quantification was determined using a 10-fold dilution series of DNA templates, derived from uninfected pennycress and pathogen-infected pennycress leaves.

**Table 2:**
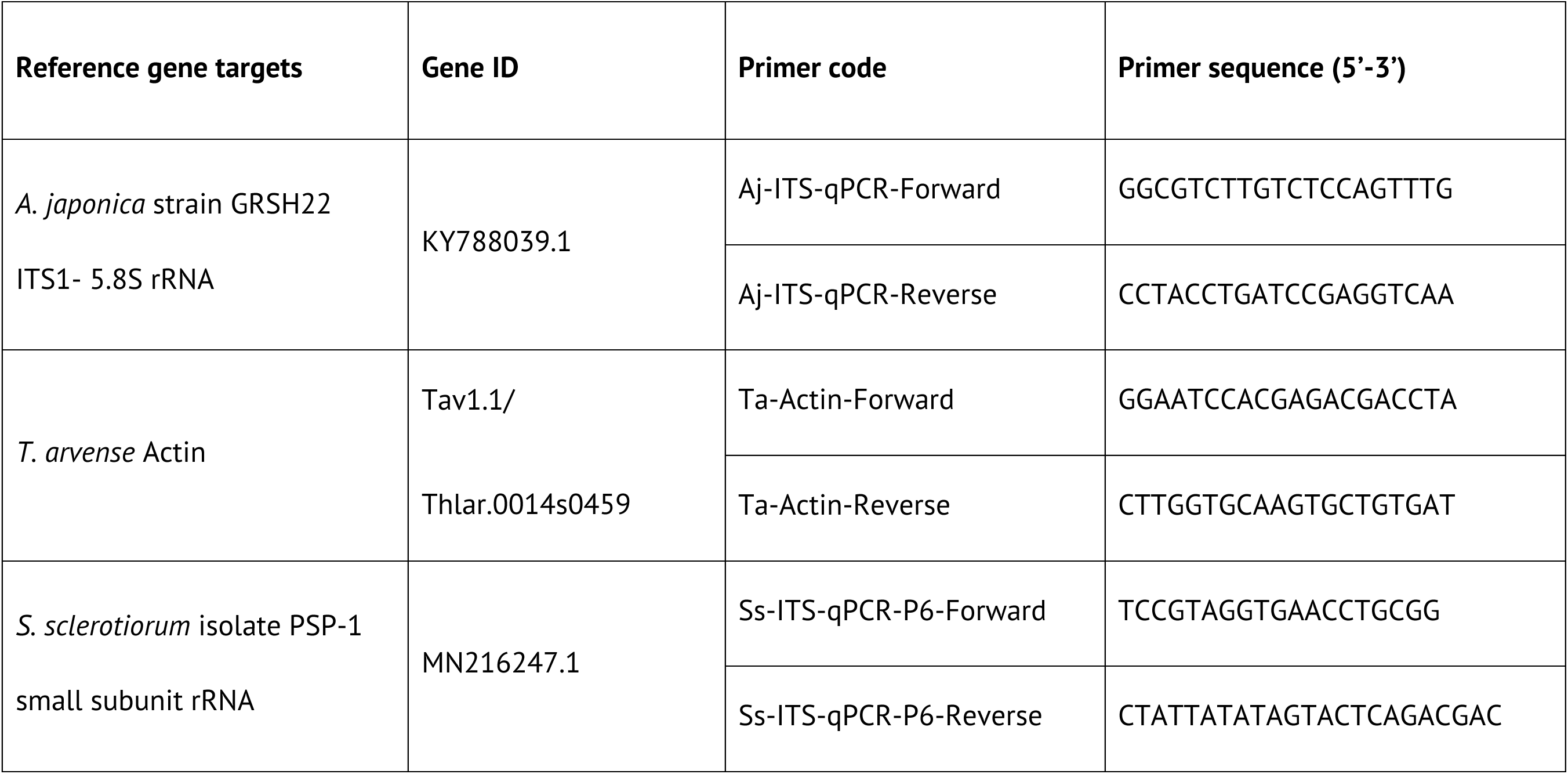
Details of primers used in this study for real-time quantification of *A. japonica* and *S. sclerotiorum* growth in pennycress

Further, qPCR primer efficiency was tested using a 10-fold dilution series of gDNA from various sources. Aj-ITS-qPCR primers used gDNA from *A. japonica* and *A. japonica-*infected pennycress leaves, while Ss-ITS-qPCR primers used gDNA from *S. sclerotiorum* and *S. sclerotiorum*-infected leaves. *Ta-*Actin primers used gDNA from uninfected and infected pennycress leaves. All primer pairs demonstrated linear amplification across a 1000-fold range of template inputs (0.05 ng/µl to 50 ng/µl) with a correlation coefficient (R²) greater than 0.95 (**Fig 5f-h**). The *TaActin* primers were 91% efficient using mock and inoculated leaf DNA as template (**Fig 5f**). The efficiency of the Ajs-ITS-qPCR primers ranged from 83-101% depending on the template (**Fig 5g**), and Ss-ITS-qPCR primers from 63-68%. We screened ten qPCR primer pairs to detect the *S. sclerotiorum* ITS but could find none with the required specificity (data not shown), which is why we proceeded with one of the selected Ss-ITS-qPCR primers despite its low efficiency (68%) (**Fig 5h**).

### DNA-based qPCR can differentially quantify *A. japonica* and *S. sclerotiorum* growth in different pennycress lines

We initially utilized the Aj-ITS-qPCR primers to detect *A. japonica* DNA on inoculated leaves and pods at 3 dpi and 2dpi, respectively (**Fig 6a-d**). Key to our approach was to isolate gDNA from roughly the same amount of tissue for each sample: a uniform 22 mm-diameter leaf disc encompassing the infected tissue, or an entire seed pod. gDNA was isolated from samples using a commercial kit (Quick-DNA Plant/Seed Miniprep Kit (Zymo Research, USA), and was diluted to 25 ng/µl for all qPCR reactions. We normalized the abundance of the *A. japonica* ITS region of the 5.8S rDNA gene to the abundance of *TaActin* using the 2^-ΔΔCt^ method. This effectively normalized the amount of *A. japonica* biomass to the amount of pennycress biomass in each sample. Using this method, the relative abundance of *A. japonica* inoculated leaves at 3 dpi was more than 1000-fold higher than mock-inoculated samples (**Fig 6b**). We observed more than 100-fold difference in relative abundance of *A. japonica* between mock-inoculated and inoculated pods (2 dpi; **Fig 6c**). We found more fungal biomass in the leaves and pods of 2032 than that of MN106 (**Fig 6a-d**), highlighting the potential of DNA-based qPCR for assessing resistance or susceptibility in pennycress lines. A non-significant difference of fungal biomass in the leaves of the two genotypes could be due to the small sample size for these experiments.

**Fig 6:**
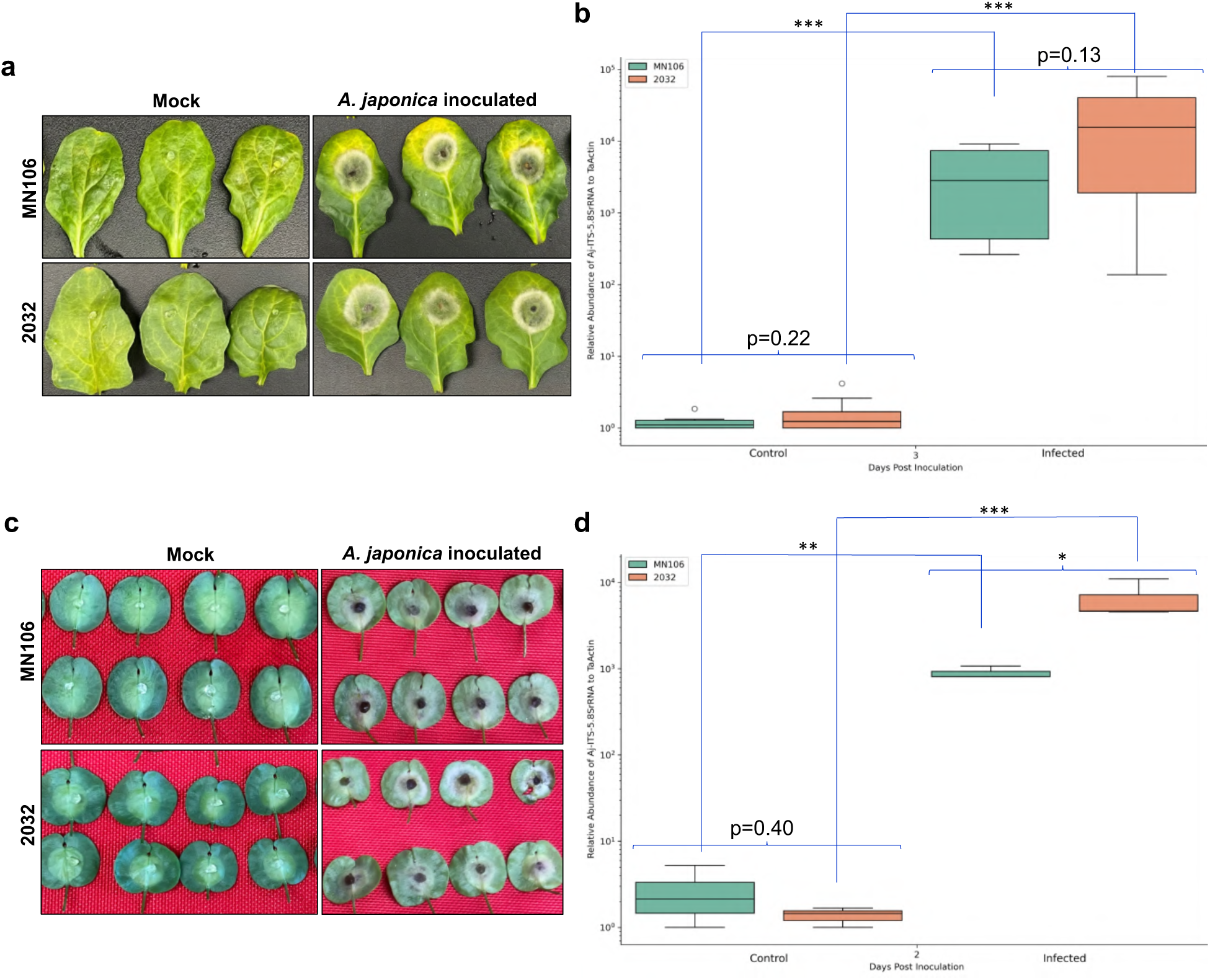
DNA-based qPCR approach for quantifying the growth rate of *A. japonica* following infection on the leaves and pods of two pennycress accessions: MN106 and 2032. Panels (a) and (c) show the phenotypes of MN106 (top row) and 2032 (bottom row), specifically their 8th leaves (a) or pods (c), in response to mock inoculation (left illustrates the *A. japonica* growth quantification for the 8th leaves of MN106 and 2032 at 3 dpi. Panel (d) represents the biomass quantification for the pods of the same accessions MN106 and 2032 at 2 dpi. The abundance of the *A. japonica* ITS 5.8S rRNA gene was normalized to the *Ta-Actin* using the 2-ΔΔCt method to determine the relative abundance of fungal biomass compared to plant biomass. Statistical significance was determined by Welch’s t-test, with a *, **, *** representing **p-value < 0.05, <0.005, and 0.0005, respectively.

To test our ability to quantify *A. japonica* over the course of an infection, we collected infected leaves and pods from MN106 and 2032 at various time points (3-6 dpi for leaves and 2-5 dpi for pods). This method showed an increase in *A. japonica* biomass on both lines over the entire time course of 3 to 6 dpi for leaves (**Fig 7a**). The increase in *A. japonica* biomass occurred from 2 to 4 dpi in pods but did not increase further from 4 to 5 dpi (**Fig 7b**). The 4 and 5 dpi time points were ideal for detecting highly significant fungal growth differences to compare MN106 to 2032 leaves (**Fig 7a**). Interestingly, we see a sigmoidal growth pattern of *A. japonica* biomass in 2032 but not in MN106 leaves. This may suggest that a more rapid growth of *A. japonica* on 2032 leaves could have exhausted all the nutrient resources by 5 dpi. A similar sigmoid growth pattern of *A. japonica* was observed on pods of both genotypes, where the growth rate plateaued from 4 to 5 dpi, but at any given time point 2-5 dpi there was signifcantly more *A. japonica* biomass on 2032 compared to MN106 pods (**Fig 7b**).

**Fig 7:**
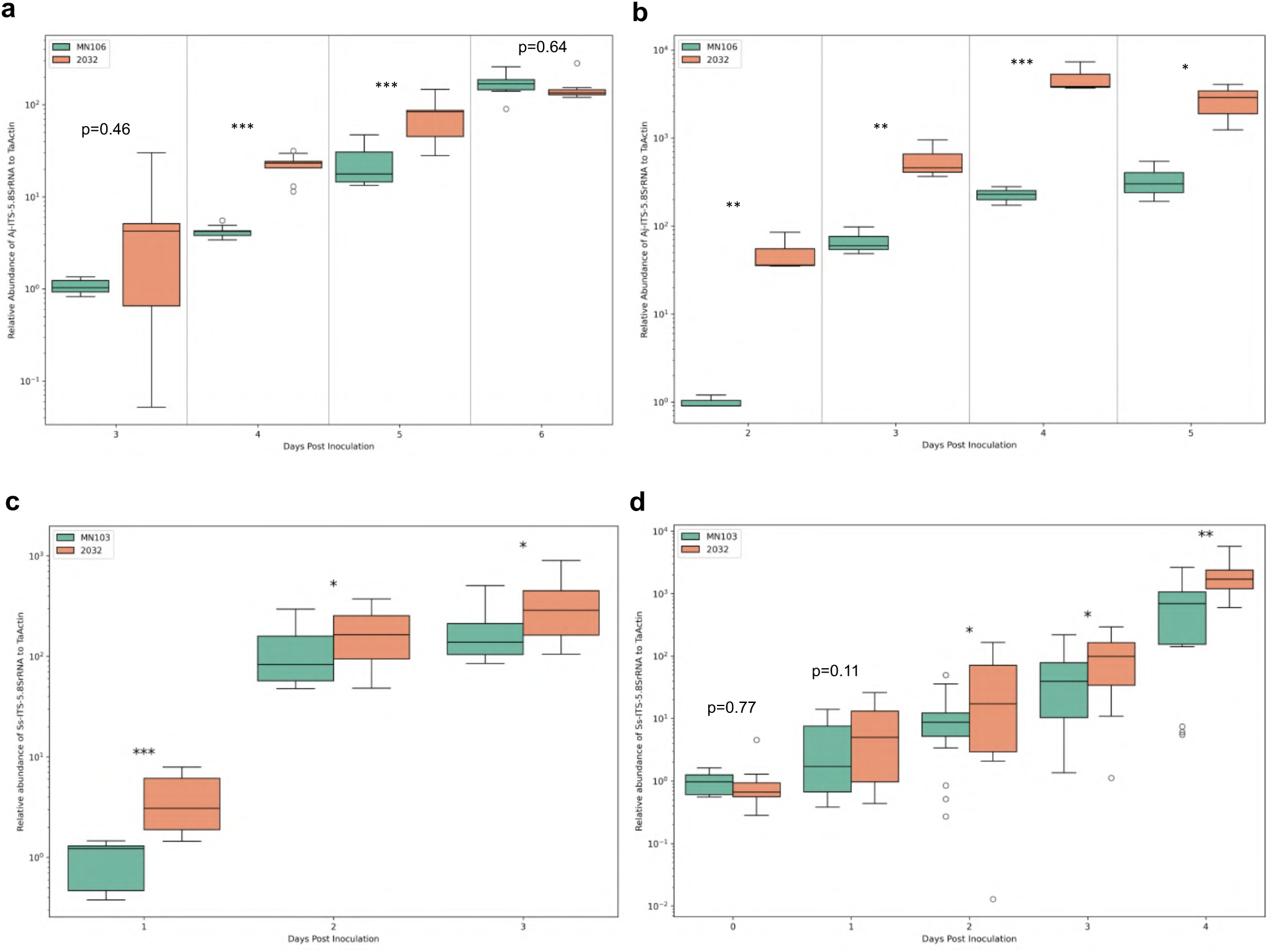
These plots illustrate the quantification of the growth rates of *A. japonica* and *S. sclerotiorum* using DNA-based qPCR during the infection process on the leaves and pods of two pennycress accessions, MN106 and 2032. Panels (a) and (b) display the qPCR-based biomass quantification of the 8th leaves (a) and pods (b), inoculated with *A. japonica* plugs over the time courses of 3-6 and 2-5 dpi respectively, for the accessions MN106 and 2032. The abundance of the *A. japonica* ITS 5.8S rRNA gene was normalized to the abundance of *Ta-Actin* using the 2-ΔΔCt method to determine the relative abundance of fungal biomass compared to plant biomass. The presented data are pooled results from two separate experiments for leaves and one for pods with each experiment containing at least three biological replicates. Panels (c) and (d) show the qPCR-based biomass quantification of *S. sclerotiorum* growth from 8th leaves, inoculated with plugs for 1-3 dpi (c) or spray-inoculated pods for 0-4 dpi (d) of accessions MN106 and 2032, using Ss-ITS-qPCR and Ta-Actin primers. The presented data are pooled results from two separate experiments for both leaves and pods. Statistical significance is denoted in terms of p-value, with *, **, and *** representing the p-value < 0.05, < 0.005, and < 0.0005, respectively based on Welch’s t-test.

Additionally, we also quantified *S. sclerotiorum* growth over time in the inoculated leaves and pods of 2032 and MN106 using the Ss-ITS-qPCR and *Ta*-Actin primer pairs by calculating the relative abundance of *S. sclerotiorum* DNA to pennycress DNA. This revealed a linear increase in relative fungal biomass from 1 to 2 dpi in leaves and from 0 to 4 dpi in pods (**Fig 7c-d**). Similar to *A. japonica* growth in pods, *S. sclerotiorum* showed a sigmoid growth pattern by plateauing at 3 dpi in leaves for both the lines, suggesting resources were starting to exhaust for the fungus (**Fig 7c**). Significantly more *S. sclerotiorum* biomass was detected on 2032 leaves compared to MN106 leaves at 1, 2, and 3 dpi (**Fig 7c**). For *S. sclerotiorum* spray-inoculated pods, there was a significant difference in fungal biomass on 2032 and MN106 pods at 2, 3, and 4 dpi but not at 0 dpi (just-sprayed) or 1 dpi (**Fig 7d**). Collectively, DNA-based qPCR effectively quantifies the growth of *A. japonica* and *S. sclerotiorum* on pennycress over time and can be used to screen for resistant and susceptible pennycress lines.

## Discussion

To the best of our knowledge, this study marks the first identification of *A. japonica* and *S. sclerotiorum* as the primary fungi responsible for black spot and white mold diseases, respectively, in pennycress—a rapidly emerging winter oilseed cash crop. Until now, there were no documented cases of pathogens causing white mold in pennycress, while *A. brassicicola* (Schwein.) being the only recorded *Alternaria* species that affects this plant (Cobb and Dillard, 1998). *A. brassicicola* is well known for causing significant damage to *Brassica* crops worldwide.

Our research introduces a rapid and precise molecular technique for diagnosing and identifying *A. japonica* and *S. sclerotiorum* from pennycress grown across various Midwest locations in North America. By utilizing advanced methods like amplification, sequencing, and homology searches of the ITS-5.8S rRNA gene, we found that *A. japonica* displays over 96% similarity, while *S. sclerotiorum* shows a perfect 100% match to strains in the NCBI GenBank database. We suggest that molecular identification of pennycress pathogens is preferred to morphological identification, particularly for *A. japonica*, whose morphology is inherently variable when culturing.

Both *A. japonica* and *S. sclerotiorum* along with other *Alternaria* species such as *A. brassicae* and *A. brassicicola* pose a substantial threat to *Brassica* crops globally (Mourou et al., 2023). These pathogens can impact plants at all stages of growth, underscoring the urgent need for effective management solutions. Breeding for resistant pennycress varieties first requires robust and repeatable assays to screen for fungal disease resistance. As such, we we developed multiple assays using field-isolated *A. japonica* and *S. sclerotiorum* strains on pennycress leaves, stems, and pods. Our findings confidently confirm the pathogenicity of both fungi on leaves and pods, with *S. sclerotiorum* also capable of infecting the stem.

Preliminary pathogenicity analyses were conducted on two natural pennycress accessions, MN106 and 2032. CoverCress Inc. had anecdotally observed that 2032 plants were highly susceptible to disease in the field and greenhouse, and in all the assays developed above we were able to demonstate that the pathogenicity of *A. japonica* and *S. sclerotiorum* was markedly higher in the 2032 accession compared to MN106.

The assessment of disease severity due to *S. sclerotiorum* was clear and easily quantifiable through visual lesion scoring, whereas the impact of *A. japonica* was less distinct. Lesions caused by *A. japonica* infection on both accessions appeared similar in contrast to the sharp, varied sizes of lesions induced by *S. sclerotiorum,* unless Trypan blue staining was employed.

Given the traditional visual-based methods to quantify resistance may not fully capture subtle differences in pathogen aggressiveness across tissues, we searched for more sensitive methods. Thankfully, the advancement in sensitive, accurate, and cost-effective DNA-based PCR assays have revolutionized pathogen diagnostics in *Arabidopsis* (Anderson and McDowell, 2015), tea (He et al., 2020), and a range of plant species, including *Brassica* crops (Mourou et al., 2023).

Anderson and McDowell (2015) and He et al. (2020) emphasized that DNA-based qPCR assays enable early-stage quantification of pathogen presence.

Building on these advancements, we have developed an affordable DNA-based qPCR assay specifically designed to quantify the growth of *A. japonica* and *S. sclerotiorum* on pennycress leaves and pods in controlled conditions. Potentially, the assay could be used to detect disease outbreaks or resistance and susceptibility in the field, but that will rely on future testing of the qPCR primers’ ability to detect the *S. sclerotiorum* and *A. japonica* strains circulating in those fields.

In our thorough investigation, we identified primers to specifically amplify the ITS-5.8S rRNA gene sequences from *A. japonica* or *S. sclerotiorum* as well as primers to amplify a housekeeping gene, *Ta-Actin* (Thalr.0014s0459) to normalize pathogen biomass against plant biomass. The choice of these genes is backed by previous studies that demonstrate the effectiveness of ITS-5.8S rRNA gene-based primers for the quick and accurate differentiation of pathogen isolates across various phylogenetic lines (Baturo-Ciesniewska et al., 2017; Blagojevic et al., 2020; Shi et al., 2021), while the *Ta-Actin* gene is recognized as a reliable reference for gene expression studies in pennycress (Chopra et al., 2018). To achieve accurate normalization, we accounted for tissue weight during gDNA extraction by harvesting infected leaf discs of the same diameter. Inoculation was carried out on detached leaves and pods, meaning that the plant tissue did not grow post-inoculation. We efficiently processed the samples using a Tissue Lyser for gDNA isolation, utilizing the budget-friendly Quick-DNA Plant/Seed Miniprep Kit, a standard method employed in many laboratories. This approach allowed for the simultaneous processing of numerous samples, significantly enhancing our capacity to manage a substantial volume of data while keeping costs low.

The qPCR assay worked well at multiple time points to assess the differences in susceptibility of MN106 and 2032. Excitingly, the qPCR assay is more sensitive than the visual based assays, especially for *S. sclerotiorum*. Significantly more *S. sclerotiorum* DNA is detected on 2032 compared to MN106 leaves at 1 dpi using qPCR, whereas it takes 2 dpi to visually see significantly larger lesions on 2032 leaves. Additionally, the qPCR is able to detect significantly more *S. sclerotiorum* DNA on 2032 pods as early as 2 dpi, prior to the development of necrotic symptoms, which otherwise first appear at 4-5 dpi. Although Trypan blue staining and qPCR both can demonstrate differences in leaf susceptibility to *A. japonica* at the same time points, the qPCR assay for *A. japonica-*inoculated pods is the only way to get a quantitative measure of susceptibility, as there were no easily measured Trypan blue lesions on pods.

Nevertheless, the qPCR assay requires multiple steps-gDNA isolation, normalization, and real-time PCR, but what makes it potentially more useful than the visual assays is that fewer inoculated samples are required. For example, significant differences in *S. sclerotiorum* lesion sizes at 2 and 3 dpi was only found in 1 out of 4 experiments, each with 6-8 plants per genotypes, necessitating pooling the data from multiple experiments. However, in a single experiment using 3 biological replicates, there was significantly more *S. sclerotiorum* DNA detected in 2032 leaves than MN106 leaves at 2 dpi (**Supplementary Fig 3**).

The future viability of pennycress as an environmentally friendly agricultural solution (a cover crop) is closely tied to our ability to tackle current challenges like fungal diseases. The techniques developed in this study not only enhance our comprehension of the complex interactions between plants and pathogens but also pave the way for identifying specific genetic mechanisms underlying disease resistance. To illustrate, we recently utilized these assays to uncover that the susceptibility of 2032 plants comes from naturally occurring mutations in the pennycress *Jumonji 14* gene (Codjoe and Kujur et al., 2025). By increasing efforts to screen for disease resistance, we can implement targeted breeding strategies and other genetic enhancement approaches that will improve the overall resilience of pennycress crops. This could lead to more sustainable agricultural practices and increased productivity in the face of environmental pressures.

## Acknowledgments

The authors thank Nathaniel Eck at the Donald Danforth Plant Science Center for his generous help with plotting the graphs and running the statistical analyses, and Jammi Prasanthi Sirasani for her help with aligning the sequences to obtain consensus sequences of *A.japonica* and *S. sclerotiorum* field isolates. We thank Daniel McLaughlin from CoverCress Inc. for sharing his expertise extracting high quality gDNA from pennycress tissues.

## Supplementary Figures

**Supplementary Fig 1:**
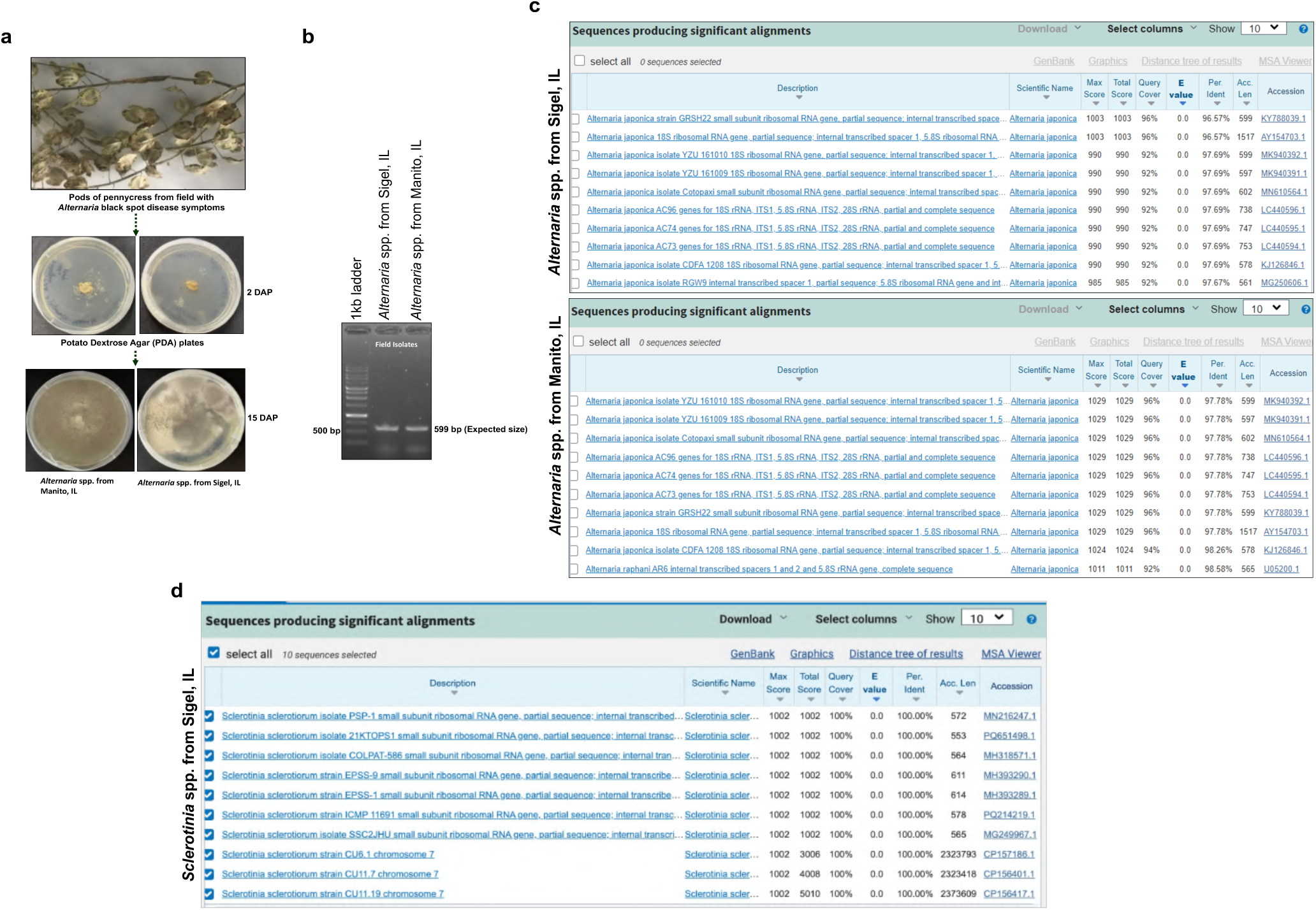
Molecular detection of *Alternaria* and *Sclerotinia* spp. field isolates collected from pods and sclerotia on stem of field pennycress was based on the amplification and sequencing of their internal transcribed spacer (ITS) region of from the 5.8S rRNA gene, obtained from field isolates of *Alternaria* spp. in Sigel, IL, and from Manito, IL, confirmed that both field isolates as *A. japonica*. d) A BLASTn search of the amplified ITS sequence from the 5.8S rRNA gene obtained from field isolate of *Sclerotinia* spp. in Sigel, IL, identified the isolate as *S. sclerotiorum*.

**Supplemental Fig 2.**
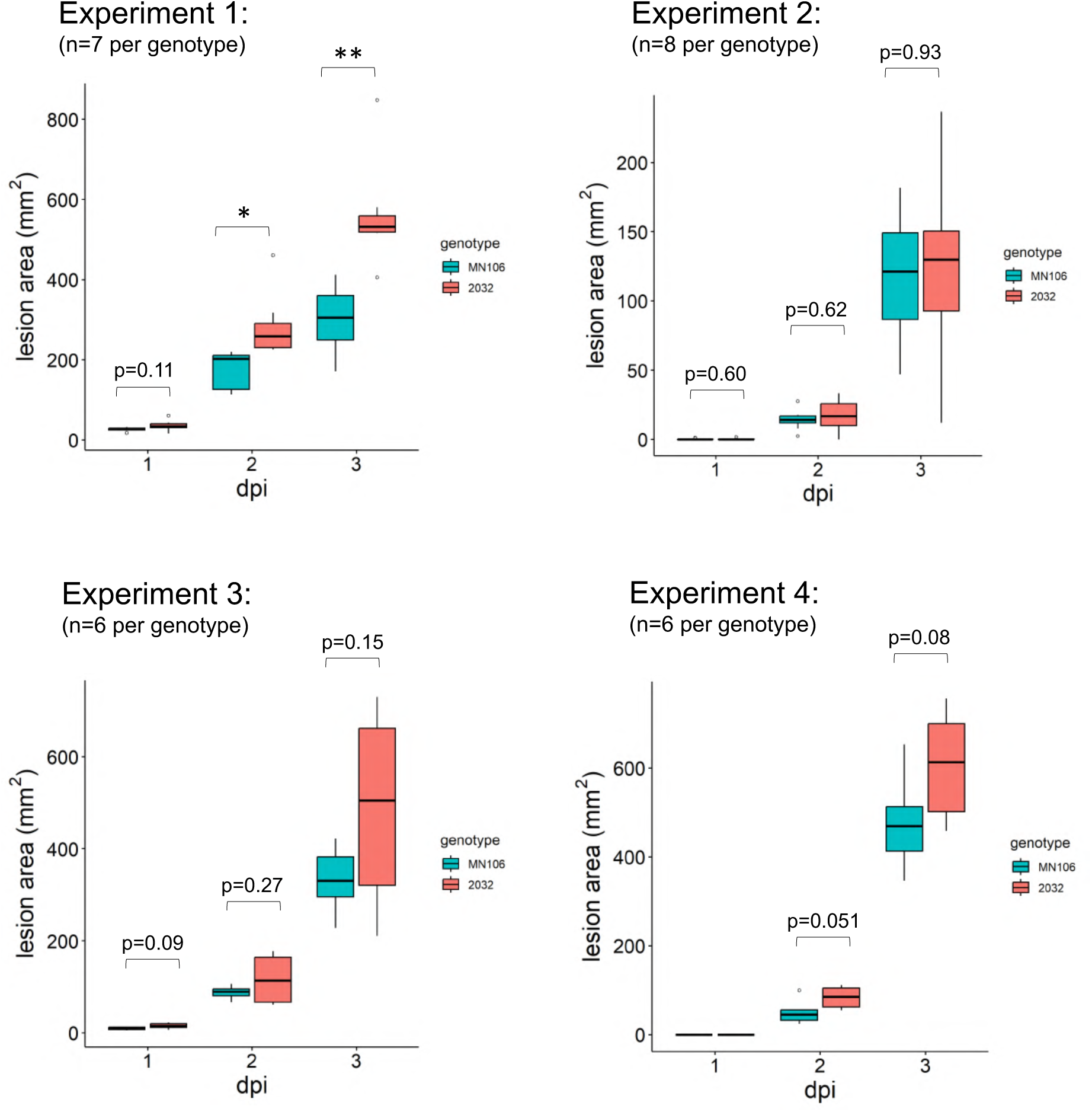
Time course analysis of *S. sclerotiorum* visual pathogenicity on MN106 and 2032 leaves across multiple experiments. The area of necrotic lesions was quantified at 1, 2, and 3 dpi. Statistical comparisons were made using Student’s t-tests. * Indicates a p-value <0.05 and ** indicates a p-value < 0.01. Data from Experiments 1, 3, and 4 were pooled together for Figure 3d. Experiment 2 was not included because the lesion sizes were much smaller than those in other experiments.

**Supplemental Fig 3.**
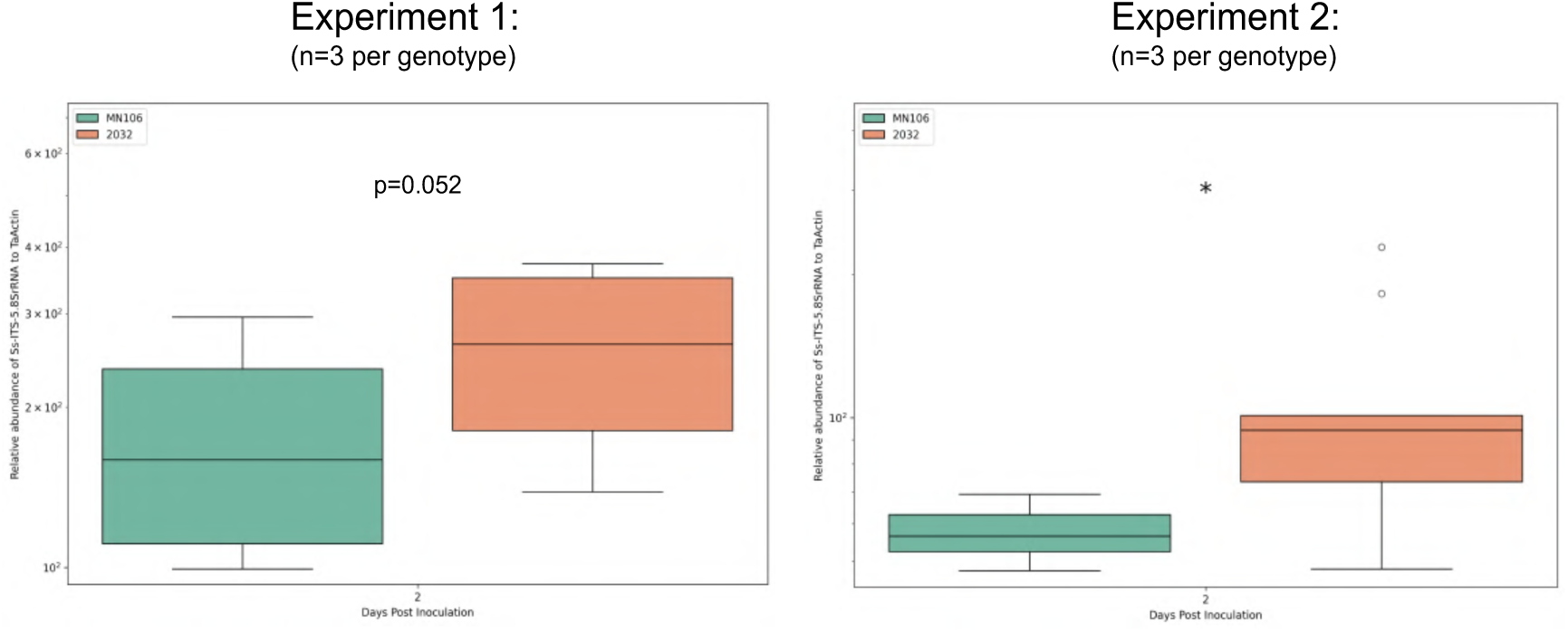
Biomass quantification of *S. sclerotiorum* using DNA-based qPCR method from the 8th leaves of two pennycress accessions, MN106 and 2032 at 2 dpi across two experiments. Each experiment was performed with 3 biological replicates, where each biological replicate had *S. sclerotiorum*-infected leaf discs from three plants. In each experiment, significantly more *S. sclerotiorum* DNA was detected in 2032 leaves than in MN106 leaves at 2 dpi. Statistical significance is denoted in terms of p-value, with *, representing the p-value < 0.05 based on Welch’s t-test.

## Literature Cited

Anderson, R. G., and McDowell, J. M. 2015. A PCR assay for the quantification of growth of the oomycete pathogen Hyaloperonospora arabidopsidis in *Arabidopsis thaliana*. Mol Plant Pathol. 16: 893–898. 10.1111/mpp.12247.

Basnet, P., and Ellison, S. 2024. Pennycress domestication and improvement efforts. Crop Sci. 64: 535–559. 10.1002/csc2.21183.

Baturo-Ciesniewska, A., Groves, C. L., Albrecht, K. A., Grau, C. R., Willis, and D. K., Smith, D. L. 2017. Molecular Identification of *Sclerotinia trifoliorum* and *Sclerotinia sclerotiorum* Isolates from the United States and Poland. Plant Dis. 101:192–199. doi: 10.1094/PDIS-06-16-0896-RE.

Blagojević, J.D., Vukojević, J.B., and Ivanović, Ž. S. 2020. Occurrence and characterization of *Alternaria* species associated with leaf spot disease in rapeseed in Serbia. Plant Pathol. 2020; 69: 883–900. 10.1111/ppa.13168

Bock, C. H., Barbedo, J. G. A., Del Ponte, E.M., Bohnenkamp, D., and Mahlein, A-K. 2020. From visual estimates to fully automated sensor-based measurements of plant disease severity: status and challenges for improving accuracy. Phytopathol Res. 2: 9. 10.1186/s42483-020-00049-8.

Chopra, R., Johnson, E. B., Emenecker, R., Cahoon, E. B., Lyons, J., Kliebenstein, D. J., Daniels, E., Dorn, K. M., Esfahanian, M., Folstad, N., Frels, K., McGinn, M., Ott, M., Gallaher, C., Altendorf, K., Berroyer, A., Ismail, B., Anderson, J. A., Wyse, D. L., Ulmasov, T., Sedbrook, J. C., and David, M, M. 2020. Identification and stacking of crucial traits required for the domestication of pennycress. Nature Food, 1: 84–91. 10.1038/s43016-019-0007-z.

Cobb, A. C., and Dillard, H. R. 1998. *Thlaspi arvense*, a new host for *Alternaria brassicicola*. Plant Dis. 82: 960960. 10.1094/PDIS.1998.82.8.960B.

Djami-Tchatchou, A. T., Tetorya, M., Godwin, J., Codjoe, J. M., Li, H., & Shah, D. M. 2023. Small cationic cysteine-rich defensin-derived antifungal peptide controls white mold in soybean. Journal of Fungi, 9:873. 10.3390/jof9090873

He, S., Chen, H., Wei, Y., An, T., and Liu, S. 2020. Development of a DNA-based real-time PCR assay for the quantification of *Colletotrichum camelliae* growth in tea (*Camellia sinensis*). Plant Methods 16: 17. 10.1186/s13007-020-00564-x

Keadle, S. B., Sykes, V. R., Sams, C. E., Yin, X., Larson, J. A., and Grant, J. F. 2023. National winter oilseeds review for potential in the US Mid-South: Pennycress, Canola, and Camelina. Agronomy J. 115: 1415–1430. 10.1002/agj2.21317

Koga, L. J., Bowen, C. R., Godoy, C. V., Oliveira, M. C., and Hartman, G. L. 2014. Mycelial compatibility and aggressiveness of Sclerotinia sclerotiorum isolates from Brazil and the United States. Pesquisa Agropecuária Brasileira. 49: 265–272.

Li, G. Q., Huang, H. C., Miao, H. J., Erickson, R. S., Jiang, D. H., and Xiao, Y. N. 2006. Biological control of *Sclerotinia* diseases of rapeseed by aerial applications of the mycoparasite *Coniothyrium minivans*. Eur J Plant Pathol. 114: 345–355. doi: 10.1007/s10658-005-2232-6.

McDowell, J. M., Hoff, T., Anderson, R. G., and Deegan, D. 2011. Propagation, storage, and assays with *Hyaloperonospora arabidopsidis*: A model oomycete pathogen of *Arabidopsis*. Methods Mol Biol. 712:137–51. doi: 10.1007/978-1-61737-998-7_12.

Meena, M., Swapnil, P., and Upadhyay, R.S. 2017. Isolation, characterization and toxicological potential of *Alternaria-*mycotoxins (TeA, AOH, and AME) in different Alternaria species from various regions of India. Sci Rep. 7: 8777. 10.1038/s41598-017-09138-9

Mourou, M., Maria, L. R., Francesco, L, and Antonia, C. 2023. Brassicaceae fungi and chromista diseases: Molecular detection and host-plant interaction. Plants 12: 1033. 10.3390/plants12173109

Sedbrook, J. C., Phippen, W. B., and Marks M. D. 2014. New approaches to facilitate rapid domestication of a wild plant to an oilseed crop: example pennycress (*Thlaspi arvense* L.). Plant Sci. 227:122–32. doi: 10.1016/j.plantsci.2014.07.008.

Shi, T., Liu, Y., Zheng, X., Hu, K., Huang, H., Liu, H., and Huang, H. 2023. Recent advances in plant disease severity assessment using convolutional neural networks. Sci Rep. 13: 2336. 10.1038/s41598-023-29230-7

Shi, X., Zeng, K., Wang, X., Liang, Z., and Wu, X. 2021. Characterization of *Alternaria* species causing leaf spot on Chinese cabbage in Shanxi province of China. J Plant Pathol 103: 283–293. 10.1007/s42161-020-00740-x

Woudenberg, J. H. C., Seidl, M. F., Groenewald, J. Z., de Vries, M., Stielow, J.B., Thomma, B. P. H. J., and Crous, P. W. 2015. *Alternaria* section *Alternaria*: Species, formae speciales or pathotypes? Studies in Mycol. 82: 1–21. 10.1016/j.simyco.2015.07.001.

Woudenberg, J. H., Groenewald, J. Z., Binder, M., and Crous, P. W. 2013. *Alternaria* redefined. Stud Mycol. 30: 75: 171–212. doi: 10.3114/sim0015.

